# Compressive Big Data Analytics: An Ensemble Meta-Algorithm for High-dimensional Multisource Datasets

**DOI:** 10.1101/2020.01.20.912485

**Authors:** Simeone Marino, Yi Zhao, Nina Zhou, Yiwang Zhou, Arthur Toga, Lu Zhao, Yingsi Jian, Yichen Yang, Yehu Chen, Qiucheng Wu, Jessica Wild, Brandon Cummings, Ivo D. Dinov

## Abstract

Health advances are contingent on continuous development of new methods and approaches to foster data driven discovery in the biomedical and clinical health sciences. Open-science offers hope for tackling some of the challenges associated with Big Data and team-based scientific discovery. Domain-independent reproducibility, area-specific replicability, curation, analysis, organization, management and sharing of health-related digital objects are critical components.

This study expands the functionality and utility of an ensemble semi-supervised machine learning technique called Compressive Big Data Analytics (CBDA). Applied to high-dimensional data, CBDA identifies salient features and key biomarkers for reliable and reproducible forecasting of binary or multinomial outcomes. The method relies on iterative subsampling, combines function optimization and statistical inference, and generates ensemble predictions of observed univariate outcomes. In this manuscript, we extend the CBDA technique by (1) efficiently handling extremely large datasets, (2) generalizing the internal and external validation steps, (3) expanding the set of base-learners for joint ensemble prediction, (4) introduce an automated selection of CBDA specifications, and (5) provide mechanisms to assess CBDA convergence, evaluate the prediction accuracy, and measure result consistency.

We validated the CBDA 2.0 technique using synthetic datasets as well as a population-wide census-like study, which grounds the mathematical models and the computational algorithm into translational health research settings. Specifically, we empirically validated the CBDA technique on a large-scale clinical study (UK Biobank), which includes imaging, cognitive, and clinical assessment data. The UK Biobank archive presents several difficult challenges related to the aggregation, harmonization, modeling, and interrogation of the information. These problems are related to the complex longitudinal structure, feature heterogeneity, multicollinearity, incongruency, and missingness, as well as violations of classical parametric assumptions that require novel health analytical approaches.

Our results showcase the scalability, efficiency and potential of CBDA to *compress* complex data into structural information leading to derived knowledge and translational action. The results of the real case-study suggest new and exciting avenues of research in the context of identifying, tracking, and treating mental health and aging-related disorders. Following open-science principles, we share the entire end-to-end protocol, source-code, and results. This facilitates independent validation, result reproducibility, and team-based collaborative discovery.

## 1. Introduction

Big Data Science is an emerging transdisciplinary field connecting the theoretical, computational, experimental, biomedical, social, environmental and economic areas. It deals with enormous amounts of complex, incongruent, and dynamic data (Big Data) from multiple sources and aims to develop algorithms, methods, tools, and services capable of ingesting such datasets and generating semi-automated decision support systems. The lack of a comprehensive mathematical formulation for Big Data Science is one of the major challenges in the development of its theoretical foundations. Other significant hurdles and gaps in Big Data pertain both the nature of the Big Data and the tools and methods to handle them. Examples of the former are Big Data heterogeneity [1], noise concentration [2], spurious correlations [3] and more. Tools and methods to handle Big Data face challenges like the choice of reliable predictive models, the specification and implementation of optimal algorithms, feasibility, scalability and convergence of the protocol on large datasets, and access to appropriate computational resources, visualization and more.

Previously, we proposed a new scalable framework for Big Data representation, high-throughput analytics (variable selection and noise reduction), and model-free inference that we called Compressive Big Data Analytics (CBDA) [4]. We showcased the robustness, efficiency, accuracy and viability of our first generation CBDA protocol on small-medium size data. In this manuscript, we expand the CBDA method and test it on large synthetic datasets (e.g., ranging from 10,000-1,000,000 cases and 1,000-10,000 features). In addition, we validate CBDA by directly applying it for detection and prediction of mood disorders (e.g., irritability) using a large population-based clinical survey. Specifically, we will validate the technique on heterogeneous and incongruent data from the UK Biobank [5, 6] datasets (see the *Datasets* section for details). The CBDA protocol relies on model-based statistical computing methods and model-free data analytics [7]. These will lead to efficient parameter estimations, reliable predictions, and robust scientific inference based on imaging, phenotypic, genetics and clinical data.

The two main strategies used by CBDA to explore the core principles of distribution-free and model-agnostic methods for scientific inference based on Big Data sets are subsampling or bootstrapping and ensemble prediction. Ensemble predictor algorithms and subsampling/bootstrapping use common approaches for objective function optimization, quantification of noise, bias, prediction error, and variance estimation during the learning/training process.

Standard ensemble methods, such as bagging and boosting [8-10], usually aggregate the results of a *single* “base” learner algorithm like support vector machine (SVM) [11] or k-nearest neighbor (kNN) [12]. CBDA employs SuperLearner [13, 14] as its ensemble predictor to combine *multiple* “base” learner algorithms into a blend of meta-learners. In addition, CBDA utilizes ensemble methods in two stages, during the Training step as well as during the subsequent Overfitting Test step (see **Fig S1** for details).

Although advanced ensemble methods like Random Forest [15, 16] could change the features’ weights during iterations, they do not directly reveal the importance of individual features. CBDA explicates the feature importance at each experimental iteration. Similar to signal estimation in compressive sensing [17], CBDA reduces the problem dimension and efficiently derives reliable and reproducible inference. In the process of computing the final inference, CBDA subsampling selects stochastically both features and cases. It identifies an optimal feature-space which may not necessarily be an average of the intermediate results.

Since its CRAN publication in 2018 [4], the CBDA package had an average of 263 downloads per months over approximately 15 months. The first version of the CBDA method was implemented as a stand-alone R package [4], which can be deployed on any desktop, laptop, or HPC cluster environment. For example, we demonstrated deploying CBDA 1.0 on a high-performance computing platform using the LONI graphical pipeline environment [18]. In this manuscript, we are enhancing the CBDA method, expanding its applications and test it on large and very heterogeneous datasets. These improvements are reflected in an integrated and upgraded CBDA 2.0 R package that is also tested on the LONI Pipeline workflow environment. In the Pipeline environment, the entire CBDA 2.0 protocol is implemented as pipeline module wrappers of various pre-processing and post-processing steps natively representing bash/shell, R, and Perl scripts that optimize the iterative CBDA subsampling phases. The latest CBDA software release is available on our GitHub repository [19].

The upgraded CBDA protocol further expands on the set of machine learning algorithms embedded in the ensemble predictor (i.e., SuperLearner [13, 14]). This new set allows the testing of several model mining performance and overall convergence metrics, which will inform the validation step and help transitioning our predictive analytics into the estimation/inference phase. The analysis of the ensemble predictor weights across the many subsamples and machine learning algorithms can also suggest a way to empirically check on CBDA computational convergence.

As a last contribution, we recast our initial mathematical formulation with the purpose of better enabling the study of the ergodic properties and the asymptotics of the specific statistical inference approaches utilized within the CBDA technique. This new simplified and more compact formulation is presented in the **Text S1.**

Our results suggest that the CBDA methodology is scalable and accurate, even with extremely high-dimensional datasets. One of the strength of combining a subsampling strategy with ensemble prediction is the ability of sifting through data where no signal is present, with low false discovery rates. The application to a real case study like the UK Biobank highlights how the protocol is flexible and scalable in a very complex predictive analytics scenario, with incongruent, heterogeneous and highly correlated data, as well as with a large degree of missingness.

## 2. Materials and Methods

This section illustrates the new CBDA protocol and methodology for representing and analyzing large datasets with binomial and continuous outcomes. First, we briefly review the main steps of the protocol and then describe in detail the new steps added to the workflow, as well as the upgrades implemented in this new study. We then review our validation procedure and results using synthetic and clinical datasets. The end-to-end CBDA processing workflow is shown in **Figure 1**.

**Figure 1:**
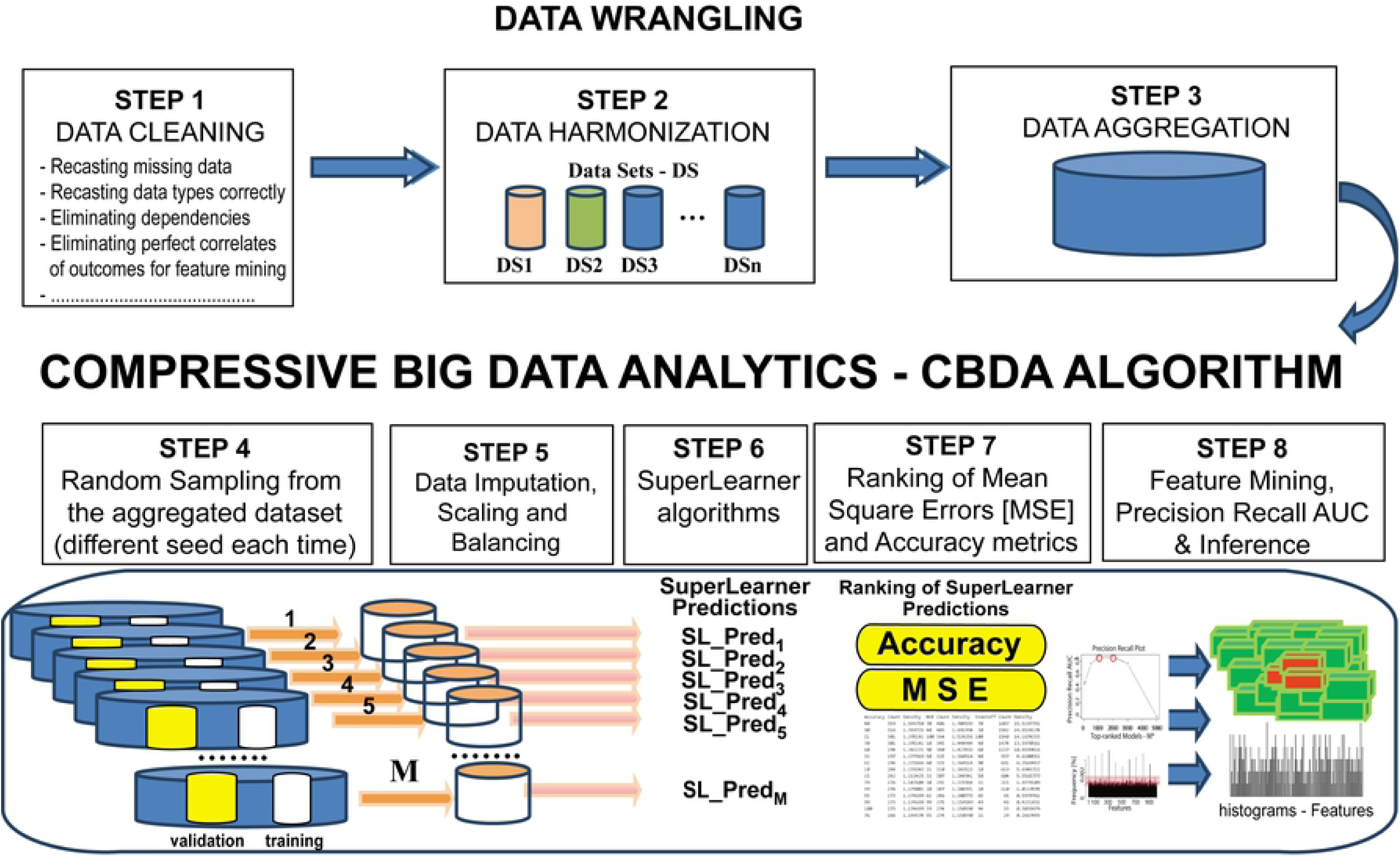
Schematic of the improved CBDA 2.0 workflow.

The entire CBDA protocol is designed, implemented and validated as a reproducible, open-project using the statistical computing language R [4, 20]. A number of training sets have been used to assess the convergence of the CBDA technique through a workflow protocol described in [4, 19]. This validation workflow runs on the LONI pipeline environment [18], a free platform for high performance computing, which allows the simultaneous submission of hundreds of independent components of the CBDA protocol (see [21] for details).

The first generation CBDA methodology and its implementation [4] can handle predictive analytics for datasets up to 1GB. Depending on the available hardware limitations, the enhanced next-generation CBDA 2.0 can handle much larger datasets. Our new implementation combines shell/bash and Perl scripts to efficiently perform data preprocessing during the subsampling steps, data staging, data post-processing, validation, and predictive analytics. The following sections outline the basics of the CBDA methodology, drawing parallels between CBDA and alternative ensemble predictor and bootstrapping strategies. Later, we will describe in detail the CBDA 2.0 implementation and highlight the new upgrades and improvements. The pseudocode is shown in the **Fig S1**.

### 2.1. A new CBDA subsampling strategy

The initial CBDA subsampling strategy is fully described in [4]. Briefly, both cases and features from the original Big Data are sampled with certain specifications given by the *Case Sampling Range* (CSR) and *Feature Sampling Range* (FSR). Our new implementation has two major upgrades in the subsampling strategy.

The main objective of subsampling is to pass a representative and balanced (if possible) but not too large sample of the Big Data to the ensemble predictor for faster analysis. In order to generalize and automate the subsample strategy, the first novelty of the CBDA 2.0 protocol is to set an upper bound in terms of number of cases and features for the subsampled datasets, namely 300 cases and 30 features. The rule of thumb here is to set the number of features as one log less than the number of cases. Other ratio cases/features can be further explored augmenting our previous report [4]). More investigation is needed to determine theoretical optimal cases/features ratio values. The goal of this study is to set a reasonably small subsample size that is sufficient to support predictive analytics and at the same time enable effective use of system resources (memory and computational cycles).

**Table 1** shows how the subsample size (e.g., 300×30), combined with the Big Data sizes, can be recasted into the previously used ranges for CSR and FSR. The new subsampling protocol significantly improves on the compression of the data needed to reconstruct the original signal (at least in the synthetic case studies) by 3-4 logs in the case of 100,000-1 million cases (see **Table 1** for details). We also reduced the total number of subsamples *M* to 5,000 (comparing to M=9,000 used in our previous study [4]). We tested CBDA 2.0 with synthetic datasets up to 70GB, but the protocol can easily be applied to larger datasets since we operate with pre-defined subsamples size (i.e., 300×30) and total number of subsamples M (e.g., 5,000). For extremely large datasets, the limiting step will be the speed in accessing a much larger dataset, which can be easily overcome by faster scripts (e.g., Python implementation).

**Table 1:** subsampling specifications for different Big Data sizes. The total number of subsamples M = 5,000.

| Description | n<br>(cases) | p<br>(features) | Sampling Rates |  |
| --- | --- | --- | --- | --- |
|  |  |  | CSR<br>(cases) | FSR<br>(features) |
| *Medium Synthetic Datasets* | 10,000 | 1,000 | 3% | 3% |
| *Large Synthetic Datasets* | 100,000 | 10,000 | 0.3% | 0.3% |
| *Very Large Synthetic Datasets* | 1,000,000 | 10000 | 0.03% | 0.3% |

Upon initializing the subsampling, the second novelty of the CBDA 2.0 protocol is a different way to perform the sampling of the validation set. In [4] we set aside 20% of the original Big Data for validation and never used it for training. However, when dealing with Big Data that approach is not scalable anymore, since the validation set might be as large as the Big Data itself. Our new strategy is then to sample a validation set each time we generate a subsample for training. Each validation set is twice the number of cases of the training set, with no case overlapping the training sample (see **Section 2.3** for details). In this study we use validations sets of 600 cases. Each sampling is done with replacement. As shown in **Table 1**, the high compression rates, achieved by fixing the max size of each subsamples, masks the potential overfitting risks. We recognize that over *M* subsamples some records might be used for both training and internal validation. However, this is only occurring in some of the *M* predictive analytics steps, and never within the same training/validation sets. Given the low CSR and FSR, the likelihood to sample the final top-ranked features in the same subsample (as resulting after the CBDA Overfitting Test stage) is almost zero.

To account for the possible multicollinearity among the set of features, we define a scaling factor that augments the FSR. We call it Variance Inflation Factor (VIF), similar to the ANOVA analysis [22].

If the subsampling is deployed on the same server location of the Big Data, the protocol performs extremely fast. The conditions are that the server has (1) a workflow system in place for the submission of multiple simultaneous jobs (e.g., LONI pipeline workflow, but it can be PBS (Portable Batch System) [23, 24], or SLURM (Simple Linux Utility for Resource Management) [25], or other schedulers), (2) shell and Perl scripting is enabled, and (3) access to an R computing environment. Our implementation uses the LONI pipeline server/client and R 3.3.3.

If any of the conditions above are not fulfilled, either the Big Data must be deployed on a server where the CBDA protocol can be executed, or, as a viable alternative, the subsampling can be done on the server hosting the Big Data and only the subsamples need to be deployed on the server where the CBDA can be executed. The latter will offer a scalable solution, since the size of the total set of subsamples is significantly smaller than the whole Big Data and can be reasonably predicted (e.g., approximately 1-2GBs).

### 2.2. CBDA ensemble prediction via the SuperLearner

The SuperLearner [13] and Targeted Maximum Likelihood Estimation (TMLE) [26, 27] theories has been developed in the past 10 years. Both are complimentary methods for parameter estimation in nonparametric statistical models for general data structures. The SuperLearner theory guides the construction of asymptotically optimal estimators of non-pathwise-differentiable parameters, e.g., prediction or density estimation, and the TMLE theory guides the construction of efficient estimators of finite dimensional pathwise-differentiable parameters, e.g., marginal means. The CBDA protocol uses SuperLearner as black-box machine learning ensemble predictor.

We can look at the SuperLearner as a data-adaptive machine learning approach to prediction and density estimation. It uses cross-validation to estimate the performance of multiple machine learning models, or the same model with different settings. The results shown in this study have been generated using 55 different classification and regression machine learning algorithms (see **Table 2**).

**Table 2:** Library of the 55 different classification and regression machine-learning algorithms used by the ensemble predictor SuperLearner in our CBDA implementation (SL.library).

| Base Learner Class | Base Learner Different specifications | Reference |
| --- | --- | --- |
| Glmnet<br>(**SL.glmnet**) | "SL.glmnet" "SL.glmnet.0" "SL.glmnet.0.25"<br>"SL.glmnet.0.50" "SL.glmnet.0.75" | [35, 36] |
| bartMachine<br>(**SL.bartMachine**) | "SL.bartMachine" "SL.bartMachine.500"<br>"SL.bartMachine.100" "SL.bartMachine.20" | [37, 38] |
| Support Vector Machine (**SL.svm**) | "SL.svm" "SL.svm.radial.10"<br>"SL.svm.radial.0.1" "SL.svm.radial.default"<br>"SL.svm.poly.2.0" "SL.svm.poly.3.0"<br>"SL.svm.poly.3.10" "SL.svm.linear"<br>"SL.svm.poly.3.n10" "SL.svm.poly.6.0"<br>"SL.svm.sigmoid" "SL.svm.poly.6.10"<br>"SL.svm.poly.6.n10" , | [11] |
| Random Forest<br>(**SL.randomForest**) | "SL.randomForest" "SL.randomForest.1000"<br>"SL.randomForest.500" "SL.randomForest.300"<br>"SL.randomForest.100" "SL.randomForest.50"<br>"SL.randomForest.20" | [15, 16] |
| Xgboost (**SL.Xgboost**) | "SL.xgboost" "SL.xgboost.500"<br>"SL.xgboost.300" "SL.xgboost.2000"<br>"SL.xgboost.1500" "SL.xgboost.d3"<br>"SL.xgboost.d5" "SL.xgboost.d6"<br>"SL.xgboost.gau" "SL.xgboost.shrink.15"<br>"SL.xgboost.shrink.2" "SL.xgboost.shrink.05"<br>"SL.xgboost.shrink.25" | [9, 10, 39] |
| K-nearest neighbor<br>(**SL.knn**) | "SL.knn", "SL.knn.100", "SL.knn.50", "SL.knn.25",<br>"SL.knn.5" | [12] |
| Others | "SL.glm" "SL.bayesglm" "SL.earth"<br>"SL.glm.interaction" "SL.ipredbagg" "SL.mean"<br>"SL.nnet" "SL.nnls" |  |

Details on default and modified parameters for each of the machine learning algorithm classes can be found in the **Text S2**. After each machine learning model has completed the analysis on each subsample, SuperLearner then creates an optimal weighted average of those models (i.e., ensemble predictor), using the test data performance. This approach has been proven to be asymptotically as accurate as the best possible prediction algorithm that is tested [13]. Although we do not directly discuss convergence results of the SuperLearner in this study, we outline a two-pronged approach for assessing the overall performance and computational convergence of our CBDA method (see **Section 2.4**).

### 2.3. CBDA two-phase bootstrapping strategy

CBDA resembles various ensemble methods, like bagging and boosting algorithms, in its use of the core principle of stochastic sampling to enhance the model prediction [28]. The purpose and utilization of the derived samples is what makes CBDA unique as it implements a *two-phase bootstrapping strategy*. In *phase one*, similar to the compressive sensing approach for signal reconstruction [29], CBDA bootstrapping is initiated by the divide-and-conquer strategy, where the Big Data is sampled with replacement following some input specifications (e.g., see section 2.1 and Table 1 here and as described in [30]). In *phase two*, during the CBDA-SuperLearner calculations, additional cross-validation and bootstrapping inputs are passed to each base learner included in the meta-learner/SuperLearner. In fact, the SuperLearner uses an internal 10-fold cross-validation, which is applied to each of the base learners in the SL.library. Moreover, one of the (optional) inputs for the SuperLearner is a set of data for external validation. In this study, the external validation set is specified as twice the number of the cases of the training chunks (600 vs. 300) and the same number of features.

Technically, this approach does not represent an external validation, since it comes from the same dataset, but no case included in the internal validation sample (600×30) would also appear in the training sample (300×30). We choose a much larger external validation set simply because there is no scarcity of data and also because the increased size does not affect CPU time (it’s just the application of the ensemble predictor model trained on the 300×30 sample).

As many distinct boosting methods can be included within the meta-learner library (e.g., XGBoost), combining the power of multi-classifier boosting within a single base learner into the larger CBDA ensemble prediction enhances the method’s power by aggregating across multiple base learners. Many studies examine the asymptotic convergence of bagging, boosting and ensemble methods [31-34]. Similar approaches may be employed to validate CBDA inference in terms of upper error bounds, convergence, and reliability. We highlight the strategies we pursue in this study in **Section 2.5**.

### 2.4. Signal filtering and False Discovery Rate (FDR) calculation

Since CBDA exploits the subsampling strategy for feature mining purposes, it is important to have an assessment of the False Discovery Rates (FDR) when ranking the most informative and predictive features. We describe now a new and general procedure to filter the signal and approximate CBDA False Discovery Rates. After the CBDA learning/training stage is completed, it is critical to determine the optimal number *M* ^*^ of top-ranked predictive models to choose from the total number *M* computed. We now outline a new procedure, which comprises two steps. The first step selects *M* ^*^ top-ranked models and plots the feature frequencies emerging from the *M* ^*^ subsamples. Then as a second step, we define a probabilistic cut-off *α* to select the likely signal (i.e., a subset of features with significantly higher frequencies). In this context, a signal is a feature frequency significantly higher than the others.

Now, *M* ^*^ can be between 0 and *M*. If *M*^*\**^ is too low or too high, no signal can be detected (see **Figure 2A-C**). For each *M* ^*^ value chosen, we plot the resulting feature frequency distribution. For *M* ^*^ = *M*, each feature frequency follows a binomial distribution, with probability 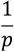. The distribution of the feature frequencies then follows a normal distribution, since it is the result of a linear transformation of a very large number of binomial stochastic variables (see [40] for details). The mean *μ* and standard deviation *σ* of the features frequency distribution across the *M* subsamples can give us a way to control the CBDA False Discovery Rate and suggests a cut-off for selecting True Positives (i.e., TP) for different *M* ^*^. By construction, if we plot the distribution of the feature frequencies across the *M* subsamples, we see an approximately normal distribution centered on the mean frequency 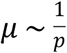 with a certain variability given by the standard deviation *σ* (see **Figure 2A**). By setting different *α* we can then control false positives (i.e., FP).

**Figure 2:**
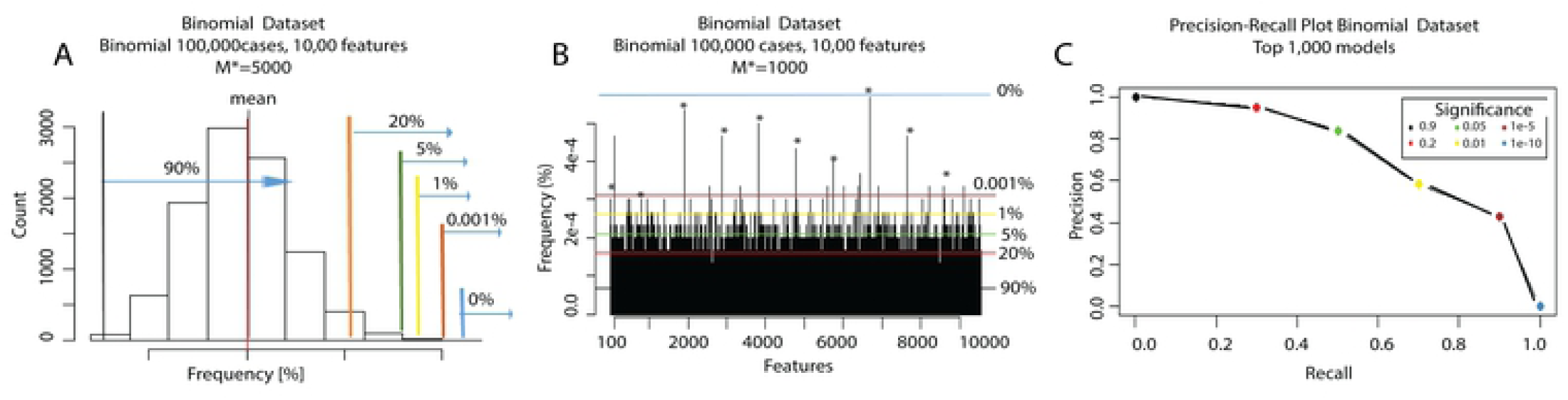
Procedure to generate Precision Recall and AUC plots. The consistent colors in the three Panels indicate identical significance levels based on the “normal” histogram in Panel A. **Panel A**: histogram of feature frequencies for the Binomial dataset with 100,000 cases and 10,000 features, for *M* ^*^ =5,000. **Panel B**: histogram of the features frequencies across the 1,000 Top Ranked models. **Panel C**: Precision-Recall plot for the Binomial dataset with 100,000 cases and 10,000 features, for *M* ^*^ = 1,000. Each circle represents a cutoff based on **Figure 2B** distribution for different *α* □. The area under the curve (AUC) in **Panel C** is then used to assess the quality and accuracy of *M* ^*^ (i.e., Top-Ranked models to consider).

For example with an *α* = *e* ^− 6^, we obtain *CBDA*_*FDR* − *cutoff*_ = *μ* + 4.75*σ*. Decreasing *α* forces a more conservative FDR. The criteria will consider any feature density value above the *CBDA*_*FDR* − *cutoff*_ as a Positive. For the synthetic datasets, the new CBDA function *CBDA_slicer()* will generate AUC (i.e., Area Under the Curve) trajectories based on different *M* ^*^ and *α*. The maximum of each trajectory will determine the optimal *M* ^*^ (see **Figure 3A-C**). The False Discovery Rate can be then calculated as the ratio of False Positives over the Positives (i.e., FP/P).

**Figure 3:**
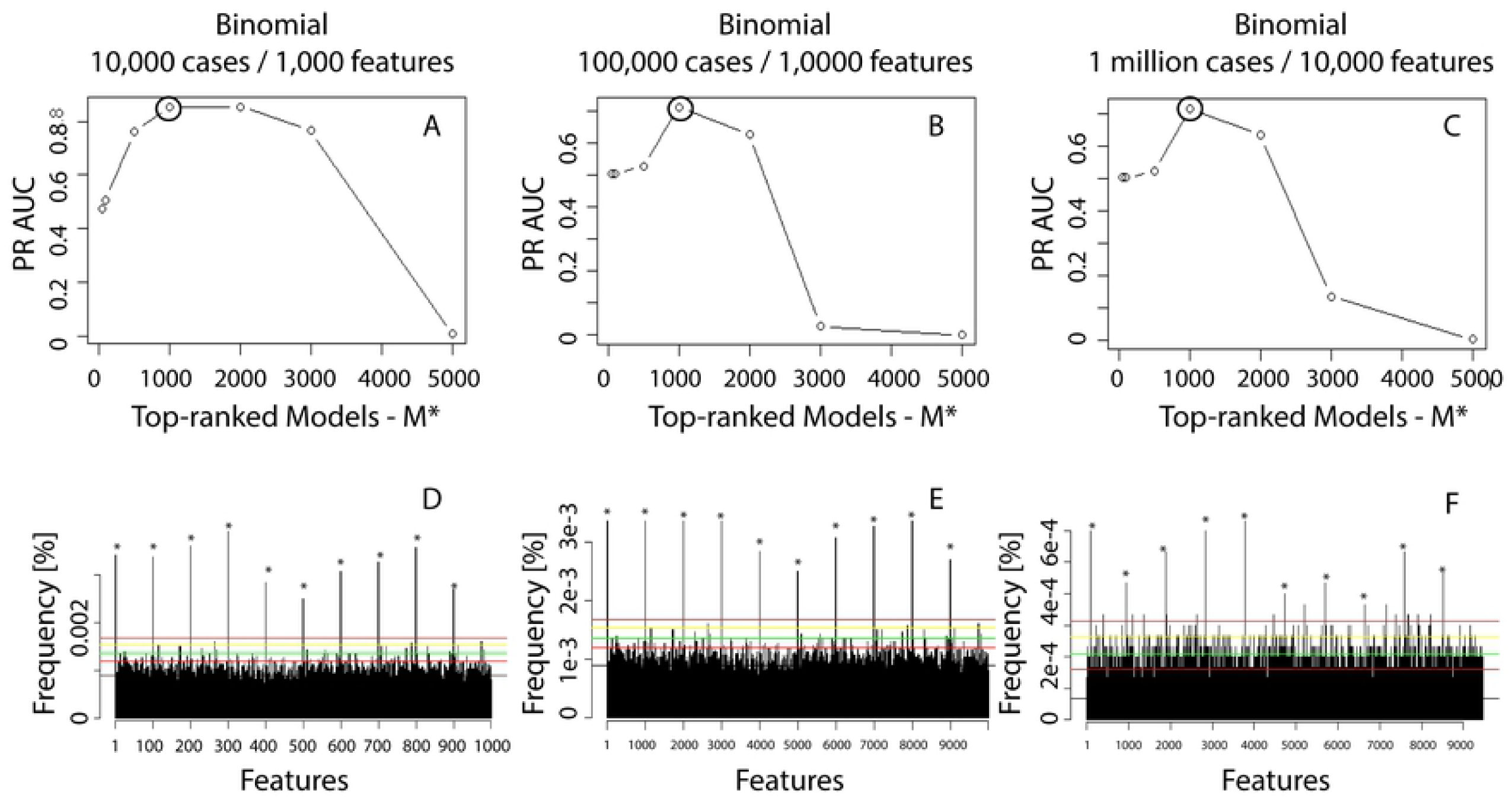
Precision Recall AUC values for different synthetic datasets and Top-Ranked models *M* ^*^. We always performed *M* = 5,000 subsamples. We calculated the PR AUV values in Panels A-C for different values of *M* ^*^, e.g., 100, 500, 1,000, 2,000, 3,000 and 5,000. The horizontal lines in Panels D-F are calculated for *α* values 0.9 (black), 1e-2 (red), 1e-5 (green), 1e-10 (yellow), 1e-16 (brown) and 1e-20 (blue). Figure 2 has more details on how the horizontal line values are calculated

**Figure 2A** shows an example of the histogram of feature frequencies resulting from one of the synthetic dataset results for *M* ^*^ =5000 (**Figure 2A**). **Figure 2** is meant to be illustrative of the methodology rather than showing the specific results. Based on the features distribution shown in **Figure 2B** and the hypothetical cut-off values set in **Figure 2A**, several *α* determine the values for the Precision-Recall (PR) plots, see **Figure 2C**, which shows the PR AUC plot for *M* ^*^ = 1,000.

In real case scenarios, the True Positive rate may not be known. Thus, the feature mining process may be guided by the optimal settings suggested by the synthetic case studies. Namely, we used the best top-ranked value (i.e., 1,000) and 5,000, 300×30 and 600×30 for the total number of samples, size of the training and validation sets, respectively.

#### 2.4.1. Precision-Recall and AUC trajectories

Usually, the number of features measured in Big Data is very large (e.g., ∼1-10K), resulting in a number of True Negatives ((TN, referring to features) that is typically 2-3 logs larger than the number of True Positives (TP). Due to this built-in class imbalance, the classical ROC curves (i.e., False Positive Rates-PPR vs True Positive Rates-TPR) are not very informative in discriminating between different *M* ^*^ decide to use Precision-Recall (PR) plots to select the best *M* ^*^ combinations. We then and *α*, since they do not account for True Negatives (TN). The definitions of Precision and Recall are given below:

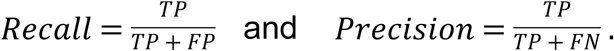

Where FP and FN represent False Positives and False Negatives, respectively. The Precision Recall Area Under Curve (PR AUC) is defined as the area under the PR curve, and that is what we use to generate our AUC trajectories. An ideal algorithm with only true positives and negatives would have a PR curve in the upper-right corner and a PR AUC of one. The new function called *CBDA_slicer()* generates a set of plots after the training stage of CBDA that display (1) the frequency of each feature as a function of the top-ranked *M* ^*^ models, (2) the correspondent PR plot, and (3) an AUC plot. The AUC plot summarizes the results of the RP as a function of *M* ^*^. In a real case scenario, the function *CBDA_slicer()* generates a final False Discovery Rate plot, instead of the AUC plot, since no true positives are known. Details on the *CBDA_slicer()* function can be found in the **Text S3** as well as in through the *help()* method part of CBDA 2.0 package [19].

### 2.5. CBDA Convergence

This section outlines two alternative approaches for testing CBDA convergence and evaluate the overall performance on a generic dataset. Both approaches rely on the analysis of the ensemble predictor output. More specifically we first look at the overall distributions of the weights/coefficients that the ensemble predictor assigns to each of the machine learning and classification algorithms throughout the training stage of the CBDA protocol (see **Section 2.5.1** for details). The working hypothesis is that if there is energy/signal/information in the data, it will be reflected in some/few algorithms being consistently more predictive than others. The assumption behind each machine learning and classification algorithm is that there is a specific correlation structure in the data, and that we can exploit it for building better predictive models. For example, when we use logistic regression models, we assume a specific relationship (e.g., log-linear) between the output to be predicted and the features (covariates or regressors) available in the dataset. Other algorithms do not return explicit relationships between outcomes and features (i.e., support vector machine, neural network, random forest), however they still superimpose specific joint distributions on the data for the purpose of improving our predictions.

The second approach examines the similarity and variability among the ensemble predictor weights/coefficients across the top predictive models. Here the assumption is that higher similarity and lower variance can suggest the convergence of the CBDA protocol (see **Section 2.5.2** for details).

#### 2.5.1. Ensemble predictor’s weights distribution analysis

The ensemble CBDA-SuperLearner model utilizes Non-Negative Least Squares (NNLS) to estimate the coefficients of a linear combination of predictive models:

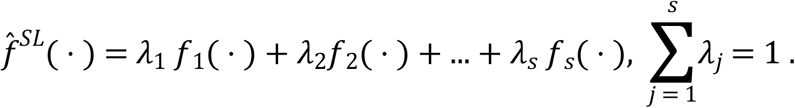

Each of the 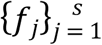 is a base learner, e.g., weak learners available in the SL.library (see **Table 2** for the list of all the base learners used in the current CBDA implementation). For each ensemble prediction model 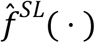, the weights/coefficients *λ*_*j*_ are optimized to return the best ensemble prediction *F*^*SL*^(*X,Y*). In other words, the NNLS step weights the performance of each algorithm in the SuperLearner training/learning step and combines them in a way that the performance of *F*^*SL*^ is as good as the best among the *f*_*j*_. Note that some algorithms might perform better than others in certain “regions” of the Big Data and by combining them, we can improve the overall meta-algorithm performance. Early studies suggested some approaches for defining asymptotic and ergodic properties of ensemble predictors [13, 27, 41, 42].

If we assume that only one or very few statistical models (e.g., priors and likelihoods function assumptions) are the “true” assumptions/models explaining the data, then only a few *λ*_*j*_ will be nonzero (or significantly non-trivial), or equivalently the set *λ*_*j*_ ≠ 0, *j* = 1,2,…,*s*, will be sparse. If we populate the SL.library with a large enough classes of learning algorithms, then a sparse set *λ*_*j*_ ≠ 0, *j* = 1,2,…,*s*, will very likely return a better prediction than a dense one.

We will investigate the relationship between the sparsity of 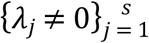 and the CBDA performance by generating enough empirical evidences to support our hypothesis. For example, we will compare the distributions of weights/coefficients between the studies on synthetic datasets generated without signal (e.g., Null datasets) and with signal, an ideal controlled scenario of white noise vs perfect information. We will then investigate the correlation between ensemble predictor accuracy or performance and the 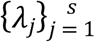 distributions using large biomedical data.

**Figures 6** and **7** show some of the results of how this approach can shed some light into the overall performance of the CBDA protocol. A new function called *SLcoef_plot()* generates a barplot of the means of the SuperLearner coefficients for each algorithm in the SuperLearner library (across the M predictive model outputs) resulting from the CBDA training stage. Details on the *SLcoef_plot()* function can be found in the **Text S3** as well as in the help of the new package [19].

**Figure 4:**
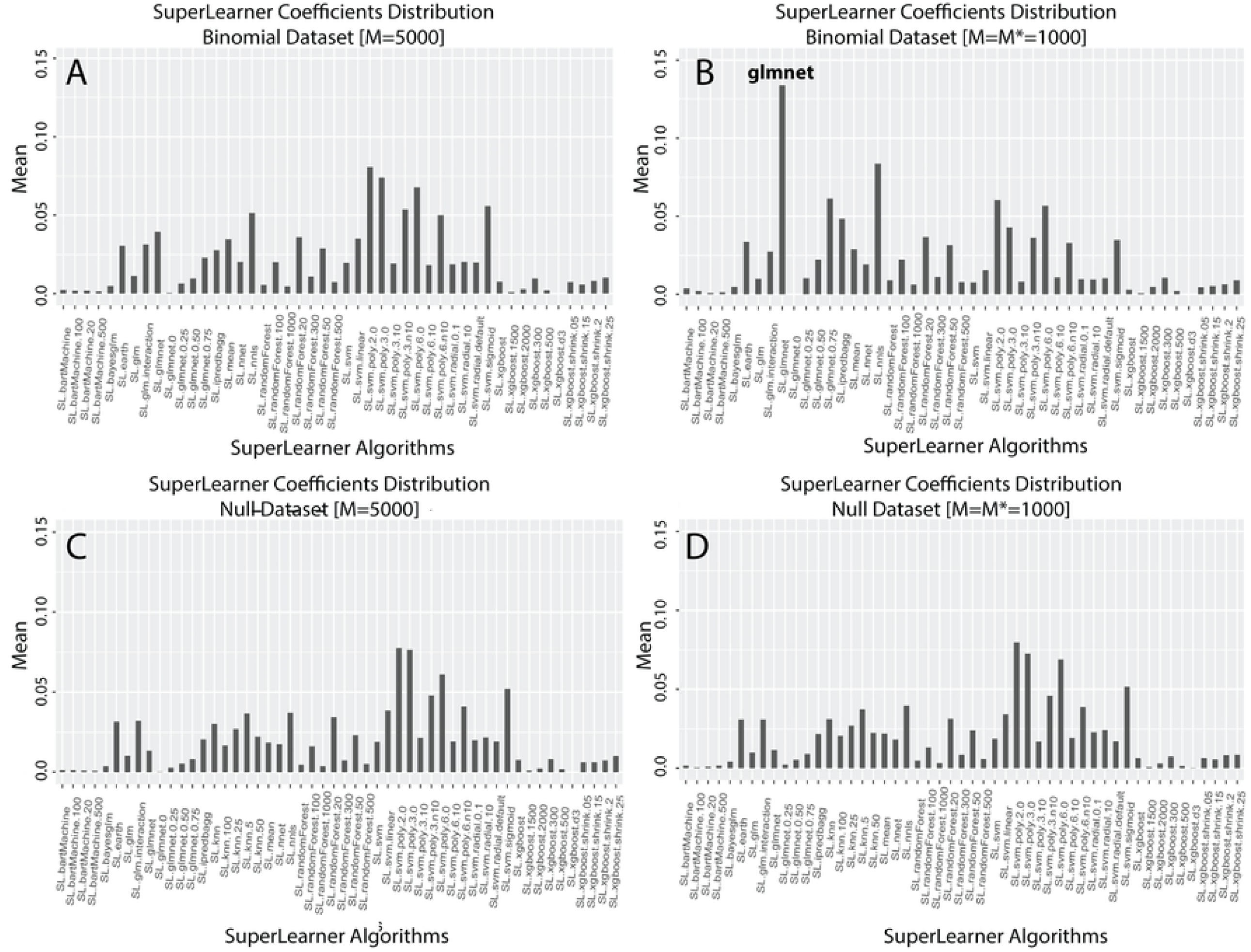
Ensemble predictor’s coefficients/weights distribution analysis, Binomial (**Panels A-B**) and Null (**Panels C-D**) datasets analysis (each with 10,000 cases and 1,000 features). The x axis displays the list of the algorithm in the SuperLearner library. The y axis shows the mean value of each SuperLearner coefficient across the *M* = 5,000 (**Panels A** and **C**) and *M* = *M* ^*^ = 1,000 (**Panels B** and **D**) top-ranked models.

**Figure 5:**
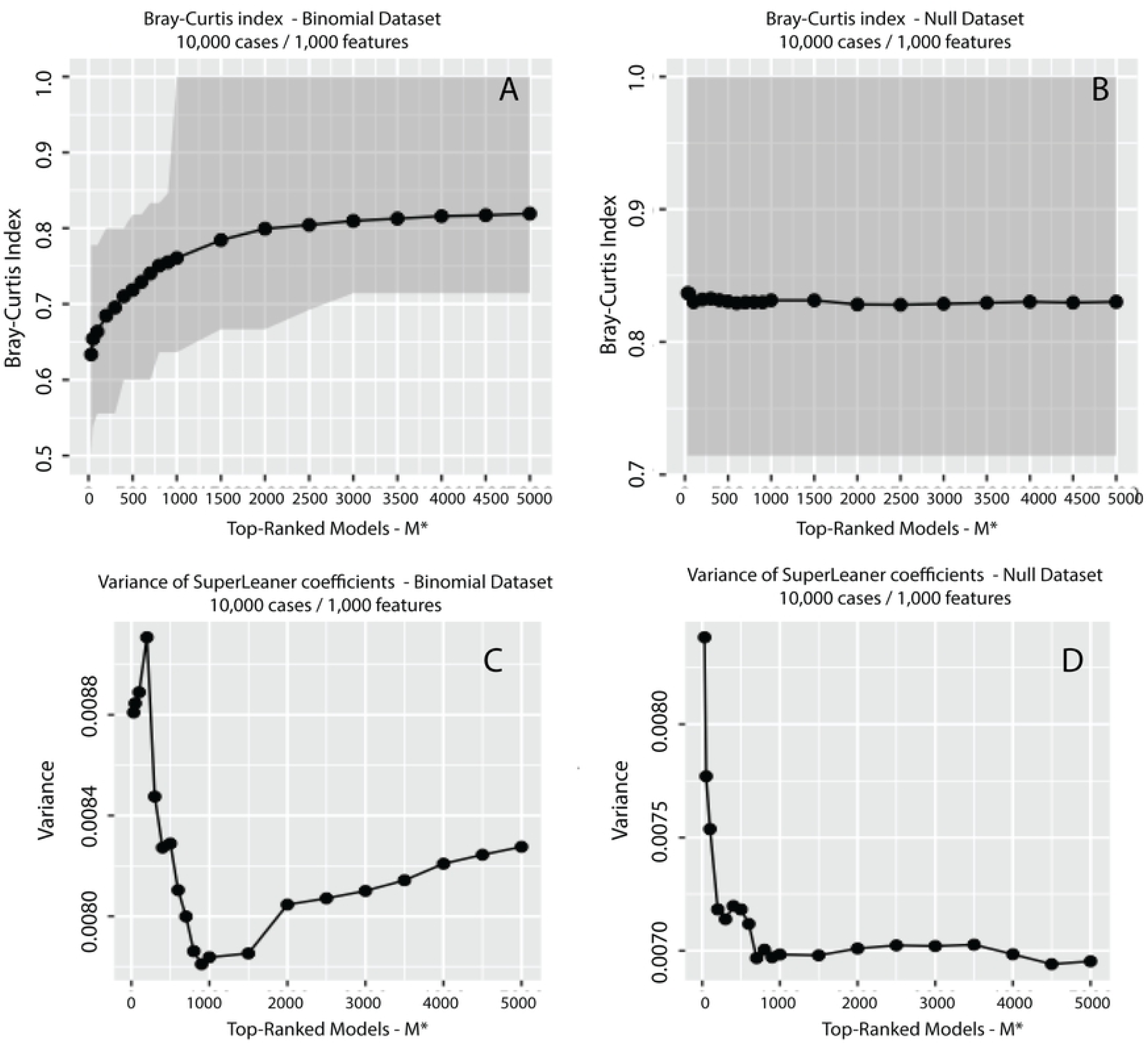
Dissimilarity and variance analysis of the coefficients/weights distributions of the ensemble predictor. Binomial (**Panels A-C**) and Null (**Panels B-D**) datasets analysis (each with 10,000 cases and 1,000 features). The x axis displays the top-ranked models *M* ^*^ (from 50 to 5,000). The y axis shows the mean value of the Bray-Curtis dissimilarity distance within the SuperLearner coefficients (**Panels A-B**) and the variance of the SuperLearner coefficients (**Panels C** and **D**).

**Figure 6:**
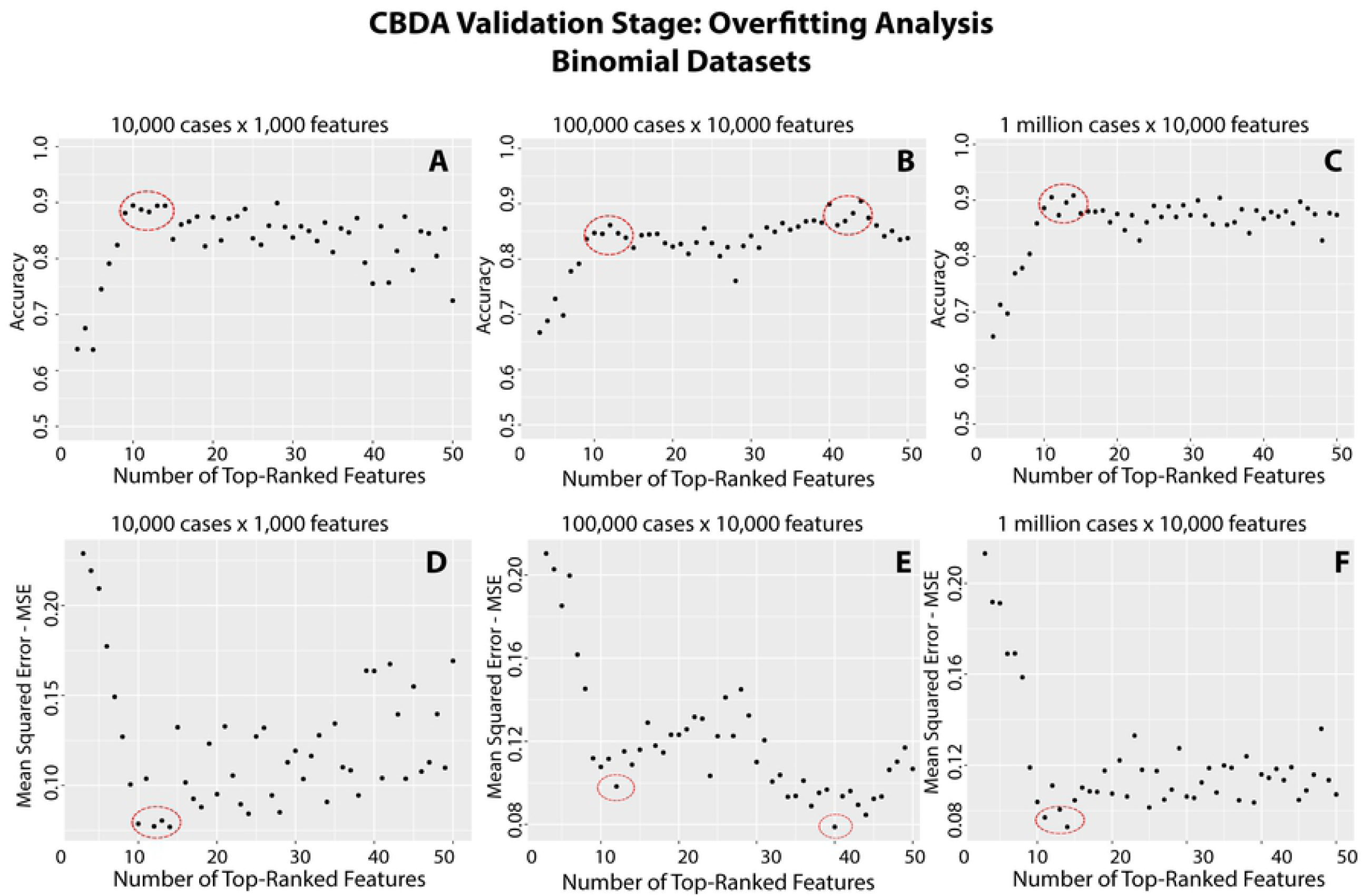
Overfitting analysis for the Binomial datasets. The y axis shows the performance metric (**Panels A-C**: Accuracy, **Panels D-F**: MSE). The x axis shows the 50 Top-ranked features resulting from the CBDA Training Stage, and used to generate the nested models during the CBDA Overfitting Test Stage on which the performance metrics are calculated. The red circles identify the likely optimal choices (i.e., max performance without overfitting) for the number of features to include in the best predictive model.

**Figure 7:**
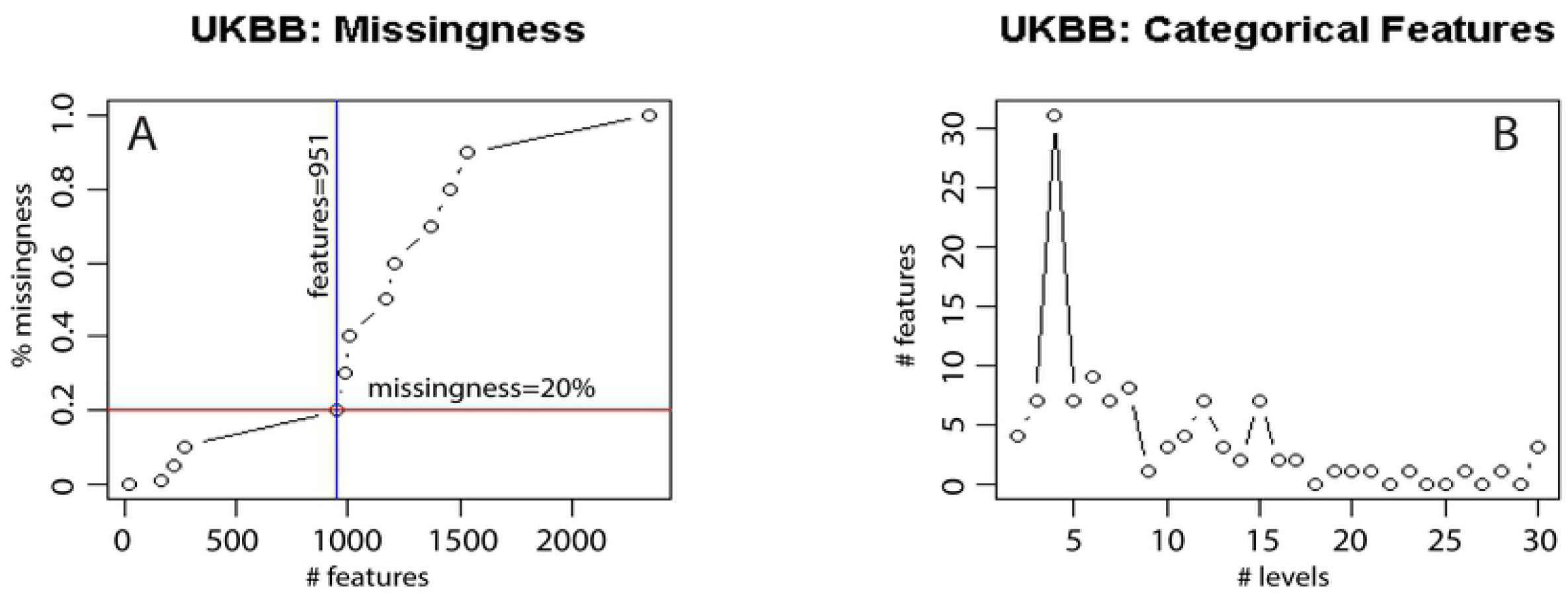
UK Biobank data wrangling statistics and results. **Panel A: Missingness analysis.** The x axis has the 2,352 features left in the physical dataset after eliminating the constant features. The y axis displays the % of missingness in the dataset, as a function of the number of features added. For example, the red horizontal line indicates a missingness of 20%, and the blue vertical line shows that if we allow for at most 20% missingness in the physical dataset, we end up with 951 features (discarding the other 1,401 features with more than 20% missingness). **Panel B**: Analysis of the 951 features left in the physical dataset. The x axis shows the number of levels (up to 30) for each of the categorical features in the physical dataset with at most 20% missingness. The goal of this analysis is to identify possible binary target outcomes for the CBDA analysis. The large peak at 4 levels shows features that are actually binary, with some incongruencies (either NAs, or values of −1 and −3 assigned to a binary outcome). The data cleaning step at this stage needs to be done manually and can only rarely be automated. **Text S4** has some detail on the list of features up to 10 unique levels.

#### 2.5.2. Dissimilarity analysis of the ensemble predictor weights

The convergence performance of ensemble predictors is typically assessed by evaluating the variance of the models predictions. For instance, in bagging, by simply calculating the variance of prediction results for each model, it is possible to assess whether the added model is useful or not. The classical strategy in bagging is described below,

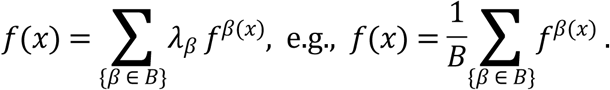

If ∑_*x*_*Var*_*β*_ (*f*^*β*(*x*)^) ≤ *α*, for some *α*, then the adding and averaging step are meaningful. In CBDA, we use a similar strategy but the ensemble model predictions are now based on subsamples of the original Big Data. CBDA does not add models, but it ranks them based on two prediction performance metrics (i.e., Mean Squared Error-MSE and Accuracy). By evaluating the similarity and variability of the weights assigned to each algorithm of the ensemble predictor across the top-ranked predictive models, we mimic the bagging strategy of tracking the variance of the model predictions to ensure convergence.

##### CBDA Implementation

After the initial steps of the CBDA protocol (random subsampling, training, ranking) of CBDA (see [4] for details), we obtain the models listed below (note: they’ve already been ranked by the accuracy/MSE):

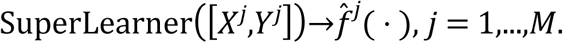

Here *M* is the total number of subsample cases, and 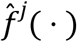 is a linear combination of the models trained by base learning algorithms in SuperLearner, which has the form of

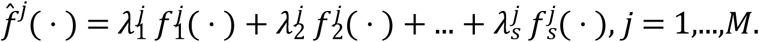

We denote the weights of the base learners as a vector 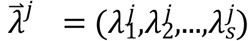 equivalent to the weights matrix:

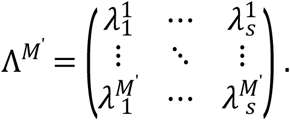

Here *M*’is a changing parameter, which ranges from the total number of subsample experiments *M*, down to a small number. In detail, we design an algorithm calculating the change of similarity among coefficient matrix by using Bray-Curtis distance [43], described as 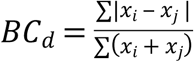. Ideally we want to see higher similarity with smaller values of *M*^’^. If the *BC*_*d*_ does not make any significant change across the decreasing values of *M*^’^, we can speculate that the performance of the CBDA protocol was not adequate. **Figure 4** shows some results of how this approach can guide our selection of the top predictive models at the end of the training stage of the CBDA protocol.

A new function called *BCplot()* generates Bray-Curtis index and variance trajectories of the M vectors of SuperLearner coefficients resulting from the CBDA training stage. Details on the *BCplot()* function can be found in the **Text S3** as well as in the help() function of the new package [19]

A new function called *Overfitting_plot()* generates two plots for each dataset after the Overfitting Test stage of the CBDA is completed (see Overfitting Plots in **Fig S1**). The x axis of each plot shows the 100 top-ranked features resulted from the CBDA training stage. The y axis shows Accuracy and MSE, respectively. By analyzing the trend of the overfitting plots, we can have a general assessment of the overall performance of the CBDA protocol on any dataset. This should be the first function to call at the end of the CBDA Overfitting Test stage. Details on the *Overfitting_plot()* function can be found in the **Text S3** and the *help()* function in the CBDA package [19].

### 2.6. Datasets

For the purpose of testing the protocol and assessing the CBDA performance. We validate the CBDA technique on three different datasets. The first two, namely the Null and Binomial datasets, are synthetically generated as cases (i.e., *n*) and features (i.e., *p*), see **Table 1** for details. Similar to our previous study, for all the Binomial datasets, only 10 features are used to generate the binary outcome variable (these are what we call truly predictive features, see details below in the Binomial Datasets section). The real case-study represents a real biomedical dataset on aging and neurodegenerative disorders (UK Biobank) [5, 6]. This data archive includes appropriate and relevant categorical (binomial/binary) outcome features, as well as clinical and neuroimaging measures.

#### 2.6.1. Null datasets

The first set of data is a “white noise” dataset (i.e., Null dataset), where the outcome *Y* is a realization of a Bernoulli vector of length *n* (i.e.,*Y* = [*Y*_1_,*Y*_2_,…,*Y*_*n*_], with *Y*_*i*_∼*Bernoulli* (0.5), *i* = 1,2,….,*n*) that is completely independent from the set of features *X*. Each column of *X* is an independent realization of a Gaussian random variable with mean 0 and standard deviation 1 (i.e., *X* = [*X*_1_,*X*_2_,…,*X*_*p*_], with *X*_*j*_ ∼ *N*(0,1),*j* = 1,2,…,*p*. We will refer to *n* as number of cases and to *p* as number of features.

### 2.6.2. Binomial datasets

The second set of data is similar to the Null dataset, but the Bernoulli vector *Y* is now an explicit function of the set of features. We establish the dependency of *Y* to *X* by selecting 10 features from *X* to build a linear additive model *Y* ∼ *X*, with non-zero coefficients for only these 10 features:

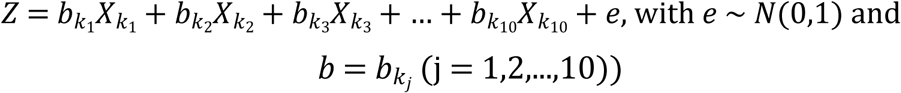

The Bernoulli outcome Y is then generated by an inverse logit on the outcome of the linear additive model (i.e., 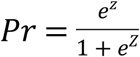 and *Y*_*i*_∼*Bernoulli*(*Pr*),*i* = 1,2,…,*n*). When necessary, various strategies may be used to binarize the predicted outcomes using the corresponding probability values.

### 2.6.3. UK Biobank archive

The UK Biobank dataset is a rich national health resource that provides census-like multisource healthcare data. The archive presents the perfect case study because of its several challenges related to aggregation and harmonization of complex data elements, feature heterogeneity, incongruency and missingness, as well as health analytics. We built on our previous UK Biobank explorative study [6] and expand to include several outcomes for classification and prediction using our CBDA protocol.

The UK Biobank dataset comprises 502,627 subjects with 4,317 features ([44] and www.ukbiobak.ac.uk). A smaller UK Biobank subset with 9,914 subjects has a complete set of neuroimaging biomarkers. By matching the ID field, we are able to merge the two datasets into a more comprehensive one with 9,914 subjects and 7,614 features (see **Table 3** for details).

**Table 3:** Aggregated **UK Biobank** clinical assessments **and** neuroimaging b**iomarker**s.

| Source | Types of Data | Sample Size |
| --- | --- | --- |

**Table 3** for details).
|  |  |  |
| --- | --- | --- |
| UK Biobank Archive<br>www.ukbiobank.ac.uk | **Baseline Characteristics:** age, DOB, sex<br>**UK Biobank Assessment Centre:** Interview information, Physical measures, Cognitive function, Imaging: MRI, DXA, Biological sampling: blood, saliva, urine<br>**Questionnaire Information:** Sociodemographics, Life style and environment, Psychosocial factors, Health and medical history, Sex-specific factors.<br>**Biological Samples:** blood, saliva, urine<br>**Genomics:** genotypes, imputation (genotypes/haplotypes), HLA<br>**Online Follow-up:** diet, cognitive function, work environment, mental health.<br>**Additional Exposure:** local environment, physical activity measurement.<br>**Health-related Outcomes:** death, cancer, stroke, myocardial infarction.<br>**Imaging Biomarkers** | The collection includes 9,914 subjects with 4,316 clinical/demographic features and 3,297 imaging biomarkers. No longitudinal data are included. The imaging biomarker data are complete, while there is a lot of missing information for the clinical/demographic features. Among the clinical/demographic features, only 23 have complete data, and 1,616 have complete missingness. |
| Challenges | ○ Build models to predict these clinical features using imaging biomarkers<br>○ Impute the missing data for the clinical/demographic features<br>○ Use text mining methods to analyze the free text. Predict clinical features using unstructured data<br>○ Validate the prediction model using the hospital admission dataset |  |

For our CBDA analysis, we then started with this subset of the entire UK Biobank dataset (i.e., 9,914 cases), with 3,297 neuroimaging biomarkers and 4,317 clinical and demographic features. Due to the high frequency of missing data in the clinical/demographic feature subset (∼72% of missingness), many of the clinical and demographic features were discarded (see **Section 3.4** for details). University of Michigan has signed a materials-transfer-agreement (MTA), 20171017_25641, with the UK Biobank consortium for the use of these UKBB data.

#### 2.6.4. Data availability

Large synthetic datasets can be downloaded using the pipeline workflow script available on our GitHub repository [45]. A client version of LONI pipeline environment [18] can be installed on a local machine and a guest account can be created with the LONI Pipeline Java/WebStart Client [46]. A pipe script can be downloaded from our GitHub repository, in the Data section [45]. After making the appropriate edits to the script in order to point to the appropriate local directories and remote data file name, the selected dataset will be compressed first and then saved in the local directory specified. Otherwise, if a client version is not available, the LONI webapp [47] can be used. Similar edits should be made to the pipeline script before loading on the LONI webapp.

Every synthetic dataset used in this manuscript can be generated from scratch using the R script in the Data Section of our GitHub repository [45]. For larger datasets, it is recommended to use the R script locally. Small size synthetic datasets are available on our GitHub repository [45]. The UK BioBank dataset is not publicly accessible, unless an IRB approval is available. A modified publicly available version of it can be downloaded from our GitHub repository [45]. A proxy of the UK BioBank dataset is publicly available on GitHub; the entire UK Biobank data is available separately (www.ukbiobank.ac.uk).

## 3. Results

We review now all the Results of the CBDA protocol applied to the synthetic datasets first, and then on the real case study datasets. We will describe the feature mining performance, test the proposed performance procedure as well as the two pronged approach in assessing CBDA convergence.

### 3.1. Binomial datasets results

The next sets of results highlight the performance of our CBDA protocol on three synthetic Binomial datasets as described in **Table 1** and in the Methods section. Each of the Binomial datasets has only 10 “true” features out of 1,000 and 10,000 features total, respectively. **Figure 3A-C** below shows the AUC trajectories based on different *M* ^*^ (from 100 up to 5,000) and *α* (from 0.9 down to *e* ^− 20^). After setting the *M* ^*^, then True and False Positives, as well as True and False Negatives are calculated for each *α* (displayed as horizontal red lines in **Figure3D-F**). **Figure 3D-F** show the correspondent frequency plot for the best *M* ^*^ (selected as the maximum of the AUC trajectories, large circles in black in **Figure 3A-C**). For each circle in **Figure 3A-C**, a histogram like **Figure 3D-F** is generated and, based on the cutoffs calculated on the histogram for *M* ^*^ = 5,000 (e.g., **Figure 2A** shows the histogram for **Figure 3B** at *M* ^*^ = 5,000), a Precision-Recall AUC curve is created (e.g., the PR AUC plot for **Figure 3B** at *M* ^*^ = 1,000 is shown in **Figure 2C**). For ease of illustration, **Figure 3** does not display the cutoff with different colors (as in **Figure 2**).

We used the accuracy of the predictions as performance metric to rank each CBDA predictive model applied to each subsample.

### 3.2. SuperLearner Coefficients Distributions: convergence by dissimilarity

Similarly to the analysis performed in **Section 3.1**, we look at the distribution of the SuperLearner coefficients/weights across the *M* ^*^ top-ranked predictive models to gain insights into CBDA overall convergence. **Figure 4** shows the results for the Binomial dataset with 10,000 cases and 1,000 features, using Accuracy as performance metric. Similar plots for all the Synthetic datasets analyzed in this study are shown in **Fig S2**, and they include both Accuracy and MSE as performance metric. For *M* ^*^ = *M* (i.e., 5,000), the coefficients/weights always show a flat distribution (**Figure 4A**), since many of the subsamples do not have any true feature in it, resulting in a poor predictive model. If we only look at the distribution of the *M* ^*^ top-ranked predictive models, some of the algorithms in the SuperLearner library performs better than others (**Figure 4B**). We could use the largest coefficient/weight obtained for *M* ^*^ = *M* as a cut-off for “true positive” algorithms in the SuperLearner library for the optimal *M* ^*^ (as returned by the procedure described in **Section 3.1**).

To validate this cut-off, we also plot the coefficients/weights distribution obtained from the analysis of the Null dataset. The same cut-off value emerges from the Null dataset analysis (**Figure 4C**). The only difference between the Null and Binomial datasets is that for the Null dataset no “better” algorithm emerges if we only look at the distribution of the *M* ^*^ top-ranked predictive models (**Figure 4D**). Thus, by comparing the SuperLearner coefficients/weights distribution between *M* and *M* ^*^, we can first check if the CBDA protocol converged to a subset of better-performing algorithms. The overall quality of the CBDA convergence can be then assessed by looking at the overall accuracy as suggested by the Overfitting Test stage of the protocol.

Another way to use the SuperLearner coefficients/weights to gain insights into the CBDA protocol convergence is to compare the Bray-Curtis (BC) dissimilarity index [43] calculated on the Binomial and Null datasets distributions, controlling for M*. **Figure 5A-B** shows the BC index as a function of *M* ^*^, from 50 to 5,000, calculated on the Binomial dataset with 10,000 cases and 1,000 features. The key information from **Figure 4** is about the dynamics of the index rather than on its values over *M* ^*^. The dynamics are unchanged in the Null case if *M* ^*^ is varied, while, in the Binomial case, the lower *M* ^*^ is, the lower the BC index is, suggesting an overall less diverse coefficients/weights distribution. The variance of the coefficients/weights follows a similar pattern when comparing the Binomial (decreasing with a minimum, **Figure 5C**) and the Null (flat, **Figure 5D**) cases. The variance also shows a result consistent with **Figure 3A**, where the variance of the coefficients/weights can be used to pinpoint the optimal *M* ^*^. In other words, we can choose *M* ^*^ as the minimum of the variance of the coefficients/weights over *M* ^*^.

### 3.3. Overfitting Test Stage

The Overfitting Test stage (see **Fig S1** for details) of the CBDA protocol generates nested nonparametric models using the top 50 (or 100) features selected in the training stage. **Figure 6** shows the performance metrics Mean Squared Error - MSE and Accuracy plotted against the number of top features used in each nested predictive model. This example uses the results obtained on the synthetic binomial datasets (as described in **Section 2.6**). The plots give us an overview of potential overfitting issues. The circles in **Figure 6** highlights the optimal number of features to include in the best predictive model, using either Accuracy or MSE as performance criteria. It is worth noting that for the Binomial datasets, the optimal number of top features to be included in the best predictive model is always consistent with the number of true features used to generate the synthetic output, i.e. 10. In fact, across the three Binomial Datasets, CBDA always ranked at least 8 of the 10 true features among the top 10. As shown by the overfitting plots in Figure 6, having 7-8 true features in the predictive model already ensures an accuracy of ∼90%.

### 3.4. Clinical Data Application: UK Biobank Dataset: data wrangling stage

We first acquire a subset of cases from the UK Biobank with complete neuroimaging measures (3,297 biomarkers) for a total of 9,914 subjects. Then we expand the data to include all the 4,316 physical features measured on these 9,914 subjects. The resulting initial merged dataset represents a second-order tensor of dimensions 9,914 × 7,614 (with one extra feature being the subject ID). The next steps were performed to ensure data harmonization and congruency.

For example, we eliminated all the constant features (1,964, all from the subset of physical features), bringing the UKBB dataset down to 2,352 physical features. Then we address the magnitude of missingness in the subset of physical features (the neuroimaging subset is complete). The plot below in **Figure 7A** shows the number of cases/subjects corresponding to different level of missingness. The more missingness we allow the more cases/subjects can be included in the final dataset. However, large missingness can seriously affect our results, no matter how efficient and accurate the imputation is. Based on the landscape shown in **Figure 7A**, we chose a 20% missingness as the best compromise that allows for a maximum number of additional features with a manageable level of missingness (increasing the missingness to 30% or 40% will only add approximately 50 subjects). Only 19 features are complete (see the list in **Text S4**).

The subset of physical features was further reduced to 951 features (still 9,914 cases). Our goal was to clean the data as much as possible before making it available to the CBDA protocol. Thus, the next data wrangling step analyzed the unique values for each feature, merging categorical and numeric (both integer and double) features. **Figure 7B** shows the number of features that have certain number of levels (or unique values), starting from 2 up to 30. We did not include the missing values code “*NA*” as a level. **Figure 7B** shows a peak at levels=4 with 31 features.

Due to the incongruency of the UKBB dataset, some obvious binary features are often coded with 4 levels (e.g., 0, 1, − 1, − 3), and the same *Field code* or ID is used multiple times for different time points. For these features, we eliminated the levels −1 and −3. The CBDA protocol will include a new module in future studies to address longitudinal data. We treated these pseudo-longitudinal data fields as the same feature and depending on the type of the measure, either exclude them or take the mean. This extra “cleaning” step reduced the physical features to 830. Unfortunately, these steps are not automated and require specific knowledge of the data under analysis, especially if the outcome of interest shows these incongruences.

Another important step is to ensure that any of the features included as “predictors” for the outcome of interest are not correlated to the outcome of interest (e.g., they measure the same or similar outcome). For example, the two features “*X2090: seen doctor for nerves, anxiety, tension or depression*” and “*X4598: ever depressed for a whole week*” are highly correlated, and using one as the outcome of interest should exclude the use of the other one as a predictor. An exhaustive and automated search will require adequate handling of unstructured data, which we will include in a separate dedicated module of the CBDA protocol. To demonstrate the CBDA protocol performance, for this study, we only chose one outcome of interest out of the 31 outcomes with 4 levels, namely “*X1940: Irritability*”. The initial levels for “*Irritability*” were − 1, − 3, 0, 1, thus we eliminated the levels 1 and 3 to make it a binary outcome The physical features labels and counts for up to 10 unique levels are shown in **Text S4**. The final UK Biobank subset analyzed to predict Irritability is 9,569 cases/subject with a total of 4,129 combined features: ID and outcome of interest (i.e., Irritability), 3,297 neuroimaging biomarkers and 830 physical features, the latter with at most 20% missing values.

### 3.5. CBDA applied to the UK Biobank Dataset

We applied the new CBDA functions to assess the CBDA performance during the training (i.e., *CBDA_slicer(), BCplot()* and *SLcoef_plot()*) and validation (*Overfitting_plot()*) stages. Ultimately the *Overfitting_plot()* results will determine the overall performance of the CBDA protocol on each dataset. **Figure 8A** shows the accuracy results of CBDA protocol for the top 100 features returned by the CBDA Overfitting Test stage executed on the neuroimaging biomarkers and the physical features as predictors. **Fig S3-A** shows the equivalent plot for the neuroimaging biomarkers only.

**Figure 8:**
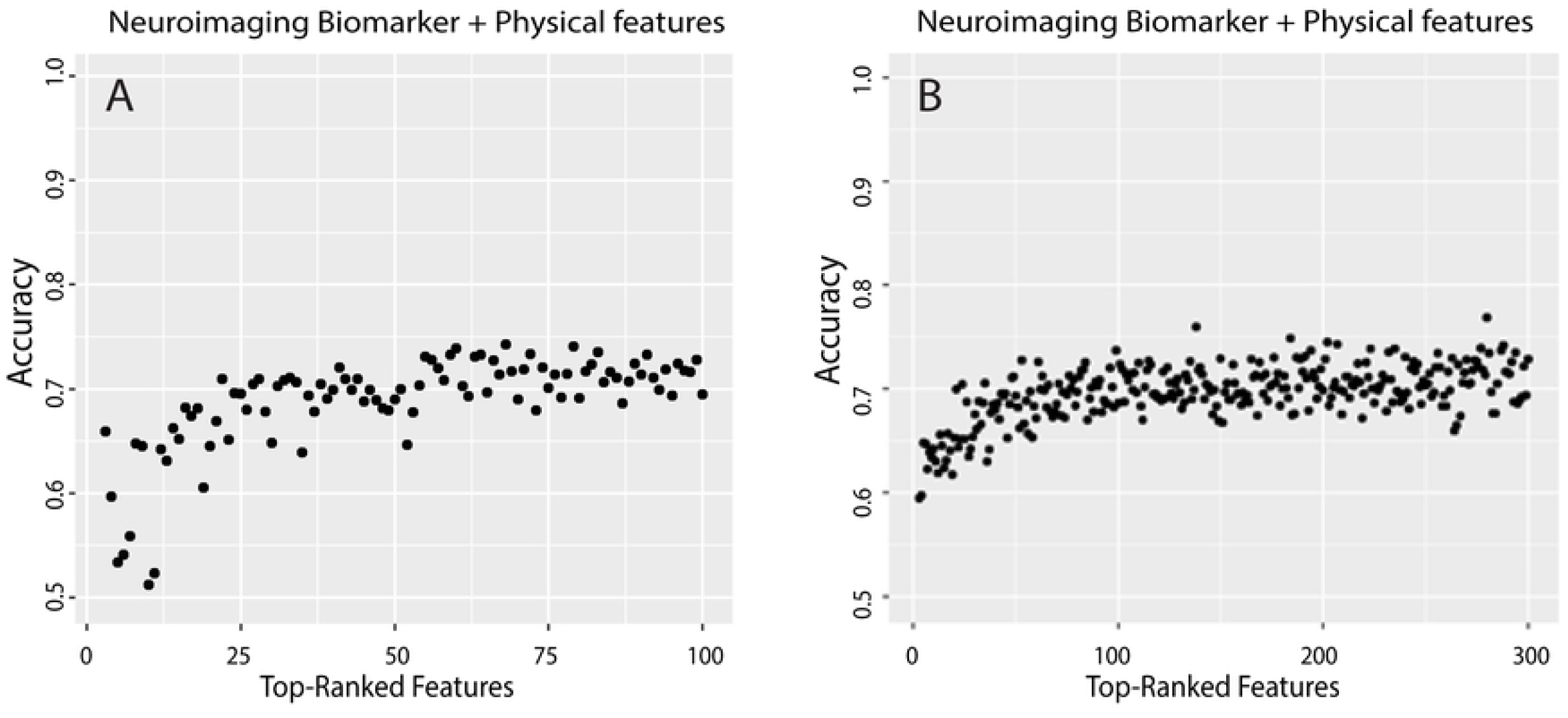
Overfitting plots for the UK Biobank dataset CBDA analysis. The x axis shows the top 100 (**Panel A**) and top 300 (**Panel B**) features returned by the CBDA training stage (see **Supplementary Figure S1** for details) on the UK Biobank dataset with both neuroimaging and clinical biomarkers. The y axis represents the accuracy of the nested models after the CBDA Overfitting Test stage. The details on the features can be found in the **Table S2** (for **Panel A**) and **S4** (for **Panel B**).

The overall accuracy converges to approximately 72% for both datasets, with a slightly different dynamic when only the top 5-10 features are included. **Tables S1** and **S2** show the lists of the top 100 features for both datasets analysis. When the physical features are included in the analysis, only 5 of them are selected in the top 100 (ranked from 20 to 24), and they all refer to neuroimaging features (i.e., see the grey cells in the **Table S2**, for details). There is a minimal overlap if we compare the top100 features, namely 4 features overlap between the 2 datasets analyses (i.e.,”rh_BA45_exvivo_thickness”, “rh_middletemporal_meancurv”, “lh_temporalpole_gauscurv”, “lh_S_circular_insula_ant_curvind”).

The highly correlated features in the biomarker dataset can explain this lack of overlap. A pairwise correlation analysis of all the neuroimaging biomarkers shows that 75% of the top 200 features (resulting after merging the 2 lists displayed in **Text S5**) have a correlation of 0.37 and higher, 50% of the top 200 features have a correlation of 0.55 and higher, 20% of the top 200 features have a correlation of 0.72 and higher, and 10% of the top 200 features have a correlation exceeding 0.82.

We ran the analysis again increasing the FSR up to approximately 180, using a *Variance Inflation Factor* (VIF) of 6. The VIF for the UKBB dataset is determined by the peculiar structure of the biomarker measures. In fact, the same measures are divided between a left and right hemisphere (factor of 2) and of approximately 15 measures on each region of interest that are correlated (e.g., volume, surface, thickness). We assumed a 20% of the number of measures for each ROI to be highly correlated and contribute to an additional scaling factor of 3 (i.e., 20% of 15). Thus, the VIF is calculated as 2 × 3 = 6, bringing the FSR from 30 to 180. Increasing the FSR further will significantly slow down the CBDA protocol without any advantages in term of performance.

The results of the CBDA analysis with the VIF=6 are shown in **Tables S3** and **S4**, where the top 300 selected features are listed for both datasets, respectively. There is an overlap of 34 features between the two analyses (see **Table 5** for details). **Fig S3-B** shows the equivalent of **Figure 8B** with neuroimaging biomarkers only included in the analysis. Increasing the FSR does not change the results in terms of overlap. If we compare the two CBDA experiments with different FSR, among the top 100 and top 300, there are 19 overlapping features for the experiment using only neuroimaging biomarkers and 9 when using both neuroimaging and clinical biomarkers. The accuracy does not change significantly over the top 50 features (although it slightly increases, see **Figure 8**). In such a highly correlated set of features, the inclusion of additional features does not affect the performance, especially if the pool of the remaining features is highly correlated. Possibly repeating the CBDA experiment a large number of times could shed some light into the correlation structure of the dataset with respect to the most predictive features, alone and in combination.

**Figure 9** shows the *SLcoef_plot()* and *BCplot()* functions output for the UK Biobank dataset with neuroimaging biomarker and physical features. The SuperLearner coefficients distribution shows how the SVM algorithm (with different kernel specifications) is consistently the most adequate to analyze the dataset (**Figure 9AB**).

**Figure 9:**
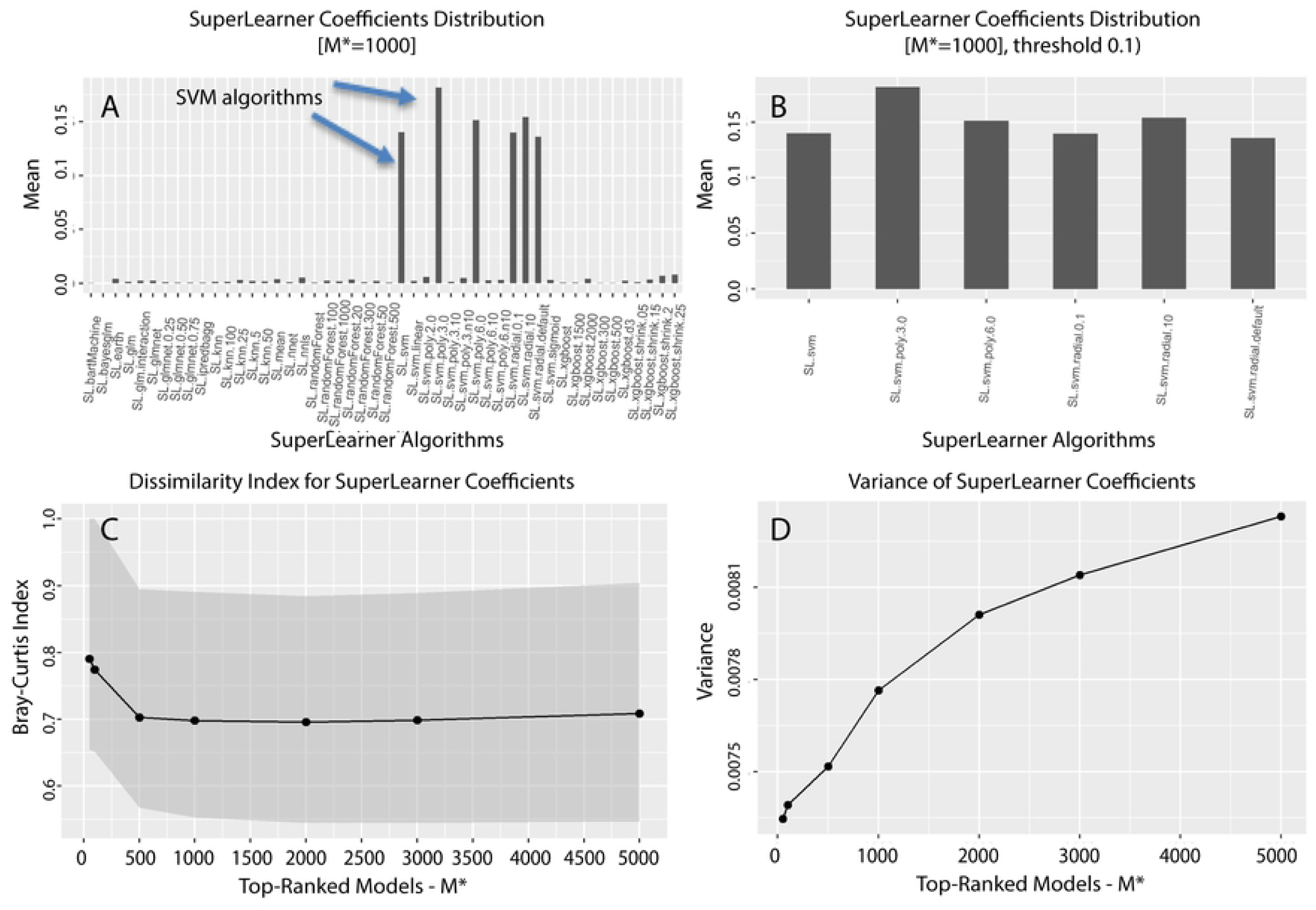
CBDA training stage for the UK Biobank dataset. SuperLearner coefficients distribution analysis (**Panels A** and **B**), **Panel C**: Bray-Curtis similarity index on the SuperLearner coefficients distribution as a function of the top-ranked models *M* ^*^. **Panel D**: variance of the SuperLearner coefficients distribution as a function of the top-ranked models *M* ^*^.

In fact, across the 55 different classification and machine learning algorithms bagged into the SL.library of our ensemble predictor, the SVM class has the best predictive power. **Figure 9A** shows the mean SuperLearner coefficients assigned during training across the 5,000 subsamples. **Figure 9B** enforces a threshold of 0.05, however most of the algorithms’ coefficients fell well below that, as shown in **Figure 9A**. No specific insights can be gained by looking at the Bray-Curtis and variance plots. The Bray-Curtis dissimilarity trajectory generated by the function *BCplot()* is relatively flat (except for an increase for *M* ^*^ < 500), with a set of minima between *M* ^*^ = 1,000 and *M* ^*^ = 3,000 (**Figure 9C**). The variance of the SuperLearner coefficients is consistently decreasing when *M* ^*^ decreases from 5,000 down to 50 (**Figure 9D**). The analysis on the UK Biobank with only the neuroimaging biomarkers returns similar results

## 4. Discussion and Conclusions

There are many challenges and opportunities associated with Big Data and team-based scientific discovery. Key components in this knowledge discovery process are curation, analysis, domain-independent reproducibility, area-specific replicability, organization, management and sharing of health-related digital objects.

Open-science offers a promising avenue to tackle some of these challenges. The FAIR data principles that we abide by (i.e., making data Findable, Accessible, Interoperable and Reusable) [48] promote maximum use of research data and foster continuous development of new methods and approaches to feed data driven discovery in the biomedical and clinical health sciences, as well as in any Big Data field.

This work expands the functionality and utility of a new method and approach that we developed in our previous study on an ensemble semi-supervised machine learning technique called Compressive Big Data Analytics (CBDA). We designed and built our CBDA protocol following the FAIR open-source/open-science principles where the scientific community can independently test, validate and expand on our second generation technology. The entire protocol, the updated R software package [4, 20] and the complete high performance computing (HPC) workflow (i.e., LONI pipeline, see [21] for details) are openly shared and publicly accessible on our GitHub repository [19]. As in our previous release, CBDA 2.0 has two open-source implementations: (1) a platform-agnostic stand-alone R package, and (2) a reproducible pipeline graphical workflow (wrapper of the R-package).

In an effort to make the CBDA protocol accessible to a larger pool of researcher so it can be deployed on virtually any HPC server, we are working now on recasting the LONI pipeline workflow into more popular and commonly used batch systems like PBS and SLURM. Currently the pre- and post-processing steps to efficiently perform subsampling are developed as shell and Perl scripts. In order to better and more efficiently handle heterogeneous, unstructured and incongruent data types, we are recasting the scripts for these two critical steps into the Python language.

Currently, our CBDA 2.0 does not handle longitudinal and unstructured data. We are developing methods and approaches to address these challenges in the context of data privacy and utility [49, 50] We will incorporate the findings and the corresponding R wrappers implementation into the CBDA protocol as soon as they are sufficiently tested and validated. A synoptic table of current and future developments for the CBDA R packages is illustrated in **Table 4**.

**Table 4:**
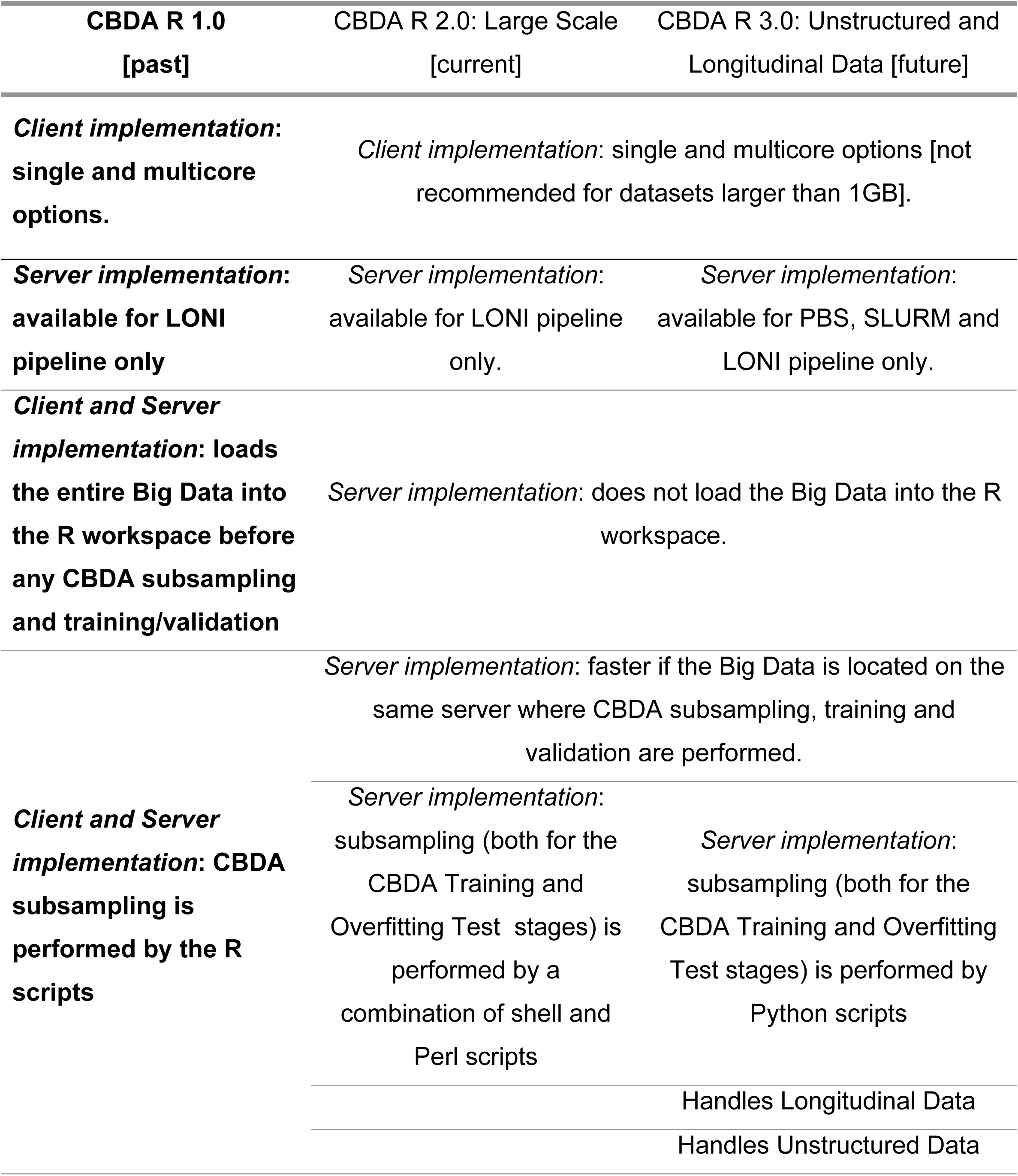
Past, present and future ***CBDA R package developments.***

We tested our second generation CBDA protocol on both synthetically generated and real datasets. Our results on synthetically generated datasets confirm and strengthen our previous study. Even with significantly reduced feature undersampling rates (e.g., from ∼1% − 5%, down to ∼0.03% − 0.3%), and increased sizes of the datasets analyzed (e.g., up to 1 million cases and 10,000 features), the CBDA protocol can identify most of the true features. Our new CBDA functionalities allows now for an immediate check on overfitting and possibly convergence issues. The new *Overfitting_plot()* function applied to our results on the synthetic datasets shows how accurate predictions can be generated even if only a subset of the true features is mined and selected.

The CBDA classification results on the UK Biobank population-wide census-like study provide empirical evidence of effective prediction, especially when the data is extremely complex, incongruent, higly correlated and has a lot of missingness. Overall, CBDA performs well with highly correlated features. Multicollinearity plays a key role in analyzing the UK Biobank. Results are not completely reproducible due to the features being extremely correlated. Also, due to the stochastic nature of the CBDA subsampling strategy, once the accuracy reaches ∼70% (soon enough, with top 10-20 features), the additional features that are added do not improve performance and are selected almost at random due to the multicollinearity. The top 10-20 features are also selected semi-randomly among the correlated top 100-300. The ranking is based on random subsampling and the feature selection in such a scenario (highly correlated features) is affected by accuracy values that are very close to each other.

The results showcase the scalability, efficiency and potential of CBDA to *compress* complex data into structural information leading to derived knowledge and translational action, where specific clinical outcomes can be targeted. Combining the CBDA and the UKBtools [51] R packages in the next wave of analysis will definitely streamline and facilitate the mapping of features, their descriptions and field ID codes, as well as the necessary data cleaning and wrangling before CBDA is implemented.

Even if this study was intended to be explorative and without a clear outcome in mind, our CBDA analysis paves the way for a deeper analysis of the UK Biobank dataset. Our results may also suggest potentially new avenues of research in the context of identifying, tracking, and treating mental health and aging-related disorders.

## Acknowledgments

Colleagues from the Statistics Online Computational Resource (SOCR), Center for Complexity and Self-management of Chronic Disease (CSCD), Big Data Discovery Science (BDDS), and the Michigan Institute for Data Science (MIDAS) provided comments and suggestions. This study was made possible and conducted under UK Biobank application number 25641.

